# Indole-3-acetic acid production is rare among gut bacteria and reflects OFOR-driven amino acid oxidation in acetogens

**DOI:** 10.64898/2026.01.29.702580

**Authors:** Mary E. DeFeo, Yuanyuan Liu, Zhiwei Zhou, Steven K. Higginbottom, Dylan Dodd

**Affiliations:** Department of Microbiology and Immunology, Stanford University School of Medicine, Stanford, CA, USA; Department of Pathology, Stanford University School of Medicine, Stanford, CA, USA

## Abstract

Indole-3-acetic acid (IAA) is a tryptophan-derived gut microbial metabolite with reported anti-inflammatory activities, but the organisms and anaerobic pathways that support robust production remain unclear. Screening 206 human gut bacterial isolates by LC-MS revealed that IAA production is rare: only five strains exceeded the limit of quantitation, and high-capacity production was confined to the acetogens *Blautia hydrogenotrophica* and *Intestinibacter bartlettii*. Across growth conditions, IAA was a minor product that rose alongside carbohydrate-sensitive, OFOR-linked catabolism of multiple amino acids, generating abundant branched-chain and aromatic organic acids. In gnotobiotic mice mono-colonized with *I. bartlettii*, these metabolites were produced *in vivo* but showed distinct host handling, with branched-chain fatty acids largely extracted between portal and peripheral plasma, whereas aromatic acids and their glycine conjugates appeared in plasma and urine. Genomic analyses and heterologous enzyme assays identified expanded repertoires of 2-oxoacid:ferredoxin oxidoreductases (OFORs) with activities spanning pyruvate/oxaloacetate, branched-chain, and aromatic 2-oxoacids, including indolepyruvate conversion to indoleacetyl-CoA, a putative intermediate *en route* to IAA. Finally, position-specific ^13^C tracing showed that CO_2_ released during amino acid oxidation is reassimilated into acetate via reductive acetogenesis, indicating that gut acetogens can maintain redox balance without fermenting partner strains. Together, these findings show that high IAA output is restricted to select gut acetogens and linked to a broader OFOR-driven anaerobic metabolism that generates additional metabolites that are absorbed by the host.

## Introduction

The gut microbiota produces a variety of small molecules that impact host physiology both locally and systemically. Bacterial tryptophan metabolites, such as indole, indole propionic acid (IPA), indole acetic acid (IAA), skatole, tryptamine, and others, circulate in human serum at similar concentrations to host metabolites^1^, and have received growing attention for their impact on human health^2,3^, particularly for their role as ligands to the aryl-hydrocarbon receptor (AhR)^4,5^. These bacterial metabolites impact organ systems across the body^6^, including immune^7,8^, gastrointestinal^9^, nervous^10^, and cardiovascular^11^, to name a few. Additionally, dysregulation in the production of these microbial metabolites is associated with diseases such as inflammatory bowel disease (IBD)^12^, multiple sclerosis^13^, and metabolic syndrome^14,15^. Therefore, understanding tryptophan metabolism in the gut microbiota could lead to a mechanistic understanding of host-microbe interactions related to health, as well as inform future therapeutics.

Indole acetic acid (IAA) is a bacterial tryptophan metabolite known for its anti-inflammatory effects, both through AhR-dependent and -independent mechanisms^16^. It is negatively associated with inflammatory conditions, such as IBD^12^ and obesity^15^, and it shows therapeutic potential in regards to colon cancer^17^ and metabolic dysfunction-associated steatotic liver disease (MASLD)^18^. While the roles of IAA in mammalian physiology are being increasingly recognized, IAA is well-established as a growth hormone in plants, commonly known as auxin^19^. Metabolic pathways for IAA production in plants and plant-associated bacteria are characterized^20^, and most of them include enzymes that require oxygen to function, such as oxidases or monooxygenases. Since the environment in the lumen of a healthy gut is anaerobic, especially in the colon where a majority of the gut microbes reside^21^, we hypothesize that gut microbes employ a different, anaerobic pathway to produce IAA.

In this study, we identify the acetogens, *Intestinibacter bartlettii* (*Iba*) and *Blautia hydrogenotrophica* (*Bhy*), as unique among >200 gut bacterial isolates in their capacity for robust IAA production. Notably, Trp-to-IAA conversion occurs as a minor component of a broader amino acid metabolic repertoire within these organisms. We implicate 2-oxoacid:ferredoxin oxidoreductases (OFORs) as key enzymes involved in branched-chain and aromatic amino acid metabolism and we provide evidence that amino acid oxidation is coupled to the Wood-Ljungdahl pathway to maintain redox balance and support acetogenesis. In vivo, these amino-acid–derived metabolites differ in systemic exposure and host processing. Together, our findings place IAA within a larger, redox-balanced metabolic program in select gut acetogens, expanding how these organisms may contribute to gut and host physiology beyond H_2_/CO_2_-dependent acetogenesis and highlighting strain- and diet-dependent levers to modulate microbiome-derived metabolite output.

## Results

### IAA production is rare among cultured isolates of human gut bacteria

To identify gut bacteria capable of producing indole-3-acetic acid (IAA), we screened a library of 206 strains spanning six common gut phyla (**Figure 1A**)^22^. Strains were grown anaerobically in carbohydrate-limited rich media supplemented with 1 mM tryptophan, and IAA in cell-free supernatants was quantified by liquid chromatography – mass spectrometry (LC-MS) after 48 hours (**Figure 1B**). Across the collection, IAA production was uncommon: only five strains exceeded the assay lower limit of quantitation (21 μM) (**Figure 1C**). All five producers belonged to the Bacillota phylum and included *Intestinibacter bartlettii* DSM 16795, two *Blautia hydrogenotrophica* strains (DSM 10507 and DSM 108220), and two *Clostridioides difficile* isolates (M68 and DSM 27543).

**Figure 1.**
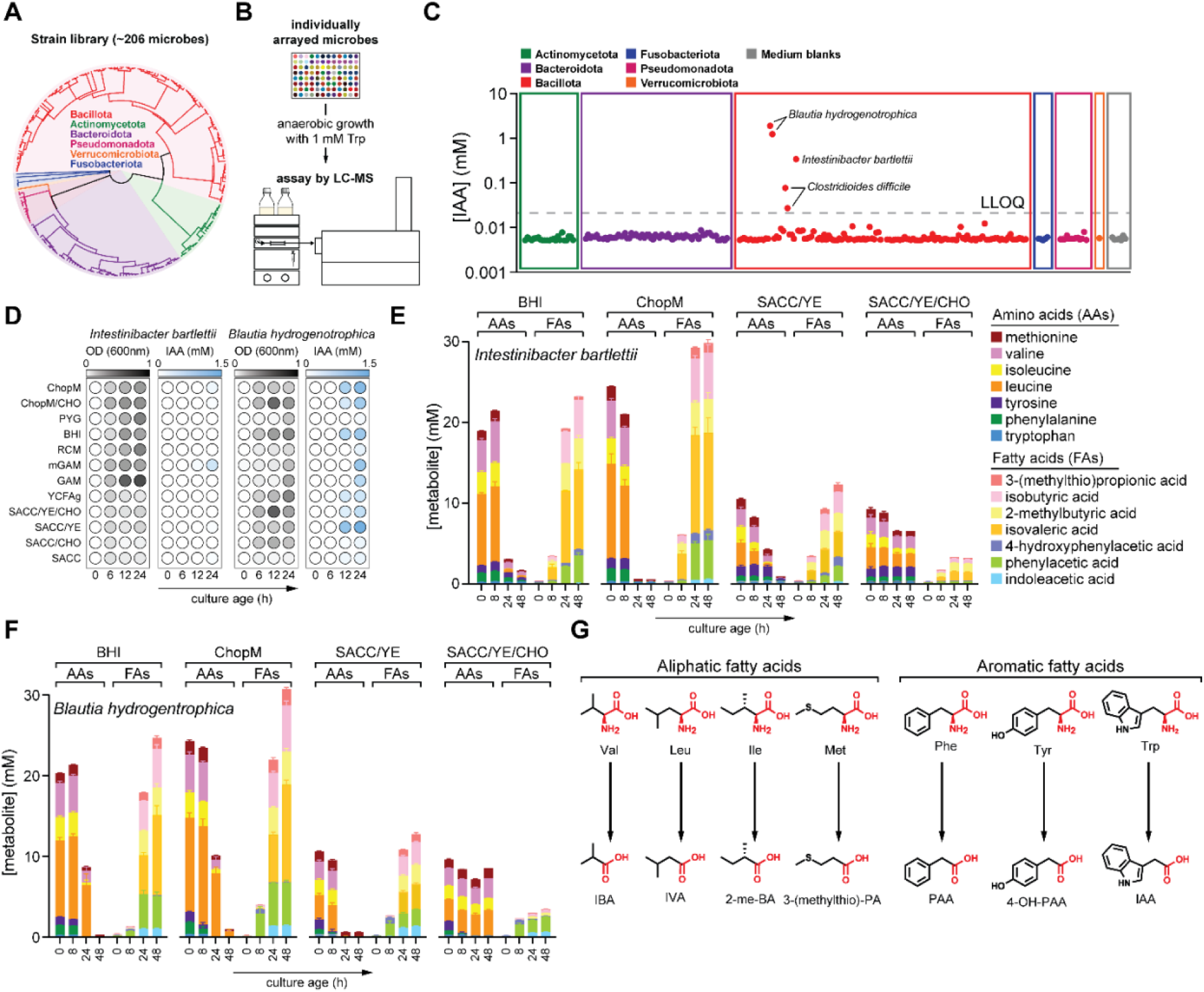
A cultivation-based screen identifies rare, high-level indole-3-acetic acid (IAA) producers among human gut isolates. (A) Phylogenetic diversity of the human gut bacterial isolate library used for screening (n = 206 strains), with major phyla indicated. (B) Overview of the screening workflow: anaerobic cultivation of isolates in microplate format followed by IAA detection by LC-MS. (C) Distribution of IAA production across all isolates. Each point represents one strain; strains are grouped by taxonomic assignment (colored blocks). The dashed line indicates the lower limit of quantitation for IAA in our assay (21 μM). (D) Growth assessed by OD_600nm_ and IAA accumulation in *Iba* and *Bhy* in 12 different media over time. (E-F) LC-MS time-course profiling of *Iba* (E) and *Bhy* (F) grown in four media, showing amino acid consumption and fatty acid production over time. (G) Schematic showing chain-shortening oxidative metabolism of multiple amino acids. For panels (E-F), data are plotted as means + SD from n = 3 replicates.

Because strains in the library were cultivated in media selected to support growth of diverse taxa, absolute IAA levels likely reflect differences in nutrient composition, including tryptophan availability in free vs. peptide form. Despite this variability, a clear hierarchy emerged: *B. hydrogenotrophica* (*Bhy*) accumulated the highest IAA concentrations (>1 mM), followed by *I. bartlettii* (*Iba*) (∼340 μM) and *C. difficile* (*Cdi*) (∼30-80 μM). A third *C. difficile* strain (ATCC BAA-1801) produced no detectable IAA under these conditions, highlighting strain-level variation.

IAA production has been reported previously for *Iba*^23^ and *Cdi*^24^. While IAA has been detected in gnotobiotic mice co-colonized with *Bacteroides thetaiotaomicron* and *Bhy*^25^, high-capacity IAA accumulation by *Bhy* in pure culture has not, to our knowledge, been reported. Although low IAA production has been described for some *Bacteroides* spp. under certain conditions^23^, none of the 66 Bacteroidota strains in our collection produced quantifiable IAA in the conditions tested here.

Together, these results indicate that IAA production is a rare phenotype among cultured human gut isolates, with *Iba* and *Bhy* emerging as high-capacity producers in our screen. We therefore used these strains as tractable models to interrogate the mechanisms that underlie IAA production.

### High-capacity IAA producers preferentially engage amino acid catabolism when carbohydrates are limited

Having identified *Iba* and *Bhy* as high-capacity IAA producers, we asked whether IAA represents a specialized, dominant metabolic output of these organisms or instead marks a broader amino acid metabolic program. From the *Bhy* strains, we focused on *Bhy* DSM 10507 because its genome sequence is available and its physiology has been studied extensively. We first examined how nutrient availability shapes growth and IAA accumulation by culturing *Iba* and *Bhy* across twelve media and sampling over time (**Figure 1D**). Two patterns were consistent across conditions: (i) *Bhy* produced higher IAA levels than *Iba*, and (ii) carbohydrate (CHO) supplementation increased growth but suppressed IAA accumulation in both strains.

Because suppression of IAA production by CHO could reflect a broader shift away from amino acid utilization, we performed targeted LC-MS profiling of amino acid and major organic acids in a subset of media (**Figure S1**). Under CHO limitation (BHI, ChopM, SACC/YE), both organisms depleted multiple amino acids and accumulated a suite of aliphatic and aromatic organic acids (**Figure 1E-F**, **Figure S1**). In contrast, CHO supplementation broadly reduced amino acid consumption and product accumulation (SACC/YE vs. SACC/YE/CHO).

Across conditions, IAA comprised only a minor fraction of the total metabolite pool relative to other fermentation end products (**Figure 1E-F**), indicating that Trp-to-IAA conversion is not the primary metabolic activity of these strains. Instead, *Iba* and *Bhy* accumulated high concentrations of products consistent with oxidation of branched-chain and aromatic amino acids, including isobutyric acid, isovaleric acid, 2-methylbutyric acid, 3-(methylthio)propanoic acid, phenylacetic acid, and 4-hydroxyphenylacetic acid (**Figure 1E-F**). Collectively, these products reached ∼30 mM in CHO-limited conditions, consistent with robust amino acid catabolism.

These metabolites resemble the oxidative branch of Stickland amino acid fermentation. However, we did not detect canonical reduced Stickland end products (5-aminovaleric acid, phenylpropionic acid, 3-(4-hydroxyphenyl)propionic acid, indolepropionic acid, or isocaproic acid)^26,27^ in either strain under the tested conditions.

Thus, in *Iba* and *Bhy*, high IAA production appears to co-occur with a broader metabolic state characterized by CHO-sensitive amino acid oxidation and accumulation of chain-shortened organic acids (**Figure 1G**).

Metabolic products of *I. bartlettii* are produced *in vivo* and enter host circulation.

We next asked whether the amino acid derived metabolites observed *in vitro* are also produced *in vivo* and whether they appear in host biofluids. Because *Bhy* colonization in gnotobiotic mice has been reported to require co-administration with other community members^25^, we focused these experiments on *Iba*.

Male (n = 5) and female (n = 3) germ-free Tac:SW mice were sampled immediately prior to gavage and two weeks after colonization. We quantified branched-chain fatty acids (BCFAs), aromatic fatty acids (AFAs), and their host-derived glycine conjugates in stool, cecal contents, peripheral and portal plasma, and urine by LC-MS (**Figure 2**). BCFAs (isobutyric, isovaleric, 2-methylbutyric acids) were near the limit of detection in stool from germ-free mice prior to gavage but were elevated in stool and cecal contents after *Iba* colonization (**Figure 2A**), consistent with *in vivo* production. For plasma, both portal and peripheral veins were sampled to determine differences in systemic availability due to hepatic processing. Free BCFAs were reduced in peripheral relative to portal plasma (**Figure 2B**), suggesting effective hepatic extraction. The corresponding acylglycines were not significantly altered by colonization in either plasma or urine (**Figure 2B-C**), consistent with limited systemic escape of microbially sourced BCFAs.

**Figure 2.**
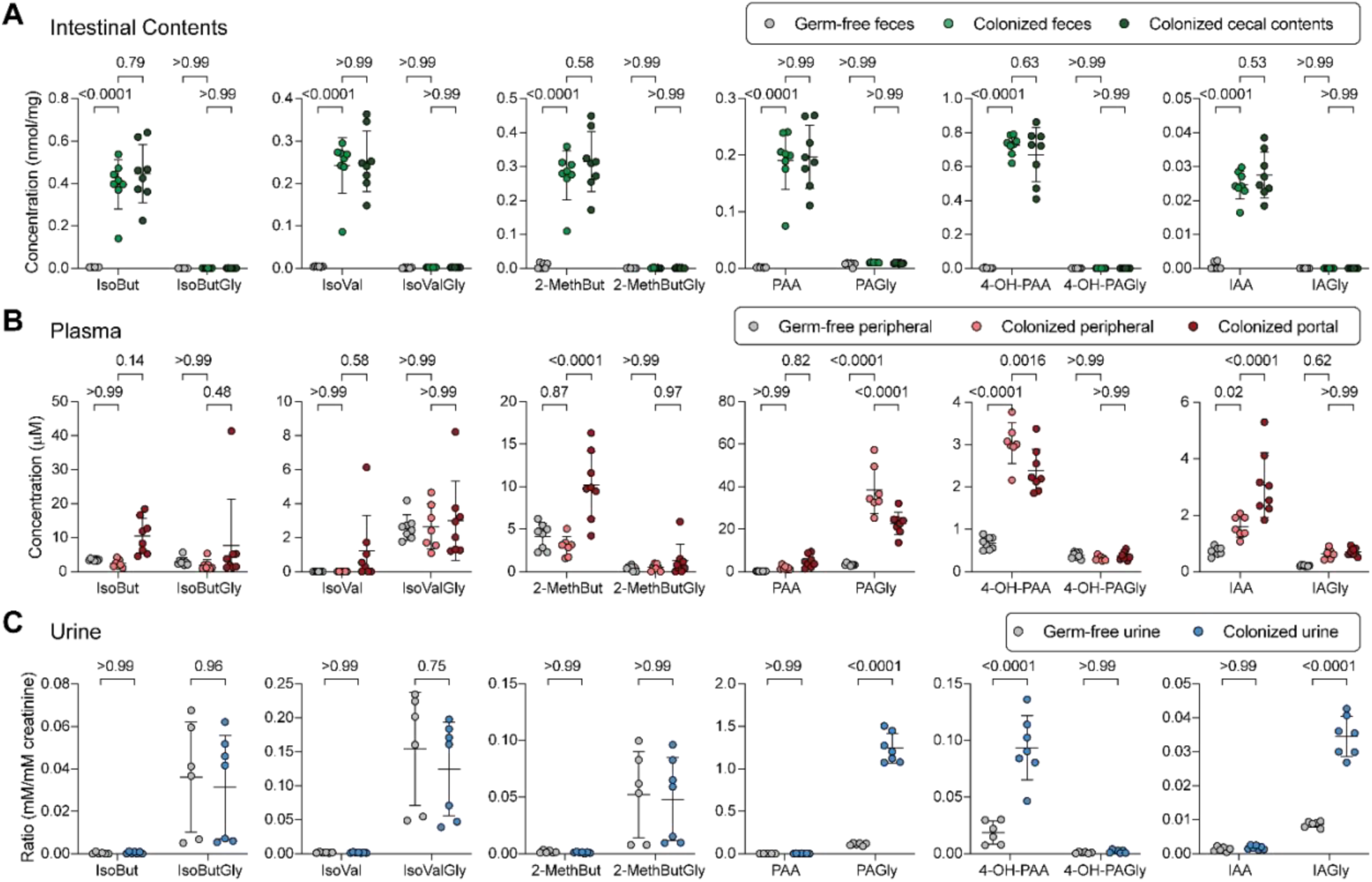
Mono-colonization with *Iba* drives compartment-specific exposure to microbial branched-chain and aromatic metabolites. Germ-free Tac:SW mice (male, n = 5; female, n = 3) were sampled as germ-free, then colonized with *Iba* and sampled two weeks later. (A) Quantification of free and glycine-conjugated branched-chain and aromatic fatty acids in feces (germ-free and colonized) or cecal contents (colonized). (B) Quantification of free and glycine-conjugated branched-chain and aromatic fatty acids in peripheral plasma (germ-free and colonized) or portal plasma (colonized). (C) Quantification of free and glycine-conjugated branched-chain and aromatic fatty acids in urine of germ-free and colonized mice. Data are plotted as individual values with lines and error bars indicating means + SD. *P*-values are from two-way ANOVA analysis.

AFAs (phenylacetic, 4-hydroxyphenylacetic, and indoleacetic acids) were also elevated in intestinal contents following colonization (**Figure 2A**). In contrast to BCFAs, AFAs and/or their glycine conjugates were increased in plasma and urine of colonized mice (**Figure 2B-C**), indicating that these microbial amino acid metabolites can reach systemic circulation and undergo host biotransformation and renal excretion. Notably, 4-hydroxyphenylacetic acid was abundant in intestinal contents despite comparatively modest production by *Iba in vitro* (**Figure 1E**), suggesting that additional variables *in vivo* (dietary substrates, intestinal transit, host processing, or enterohepatic cycling) influence steady-state levels. Phenylacetic acid circulated predominantly as phenylacetylglycine and was excreted at high levels in urine (**Figure 2C**). IAA was present in intestinal contents and plasma largely in its free form but was preferentially excreted in urine as indoleacetylglycine (**Figure 2A-C**), paralleling prior observations for indolepropionic acid metabolism in mice^27^.

Together, these results support the conclusion that *Iba* produces both BCFAs and AFAs *in vivo* and that these metabolites can be absorbed and differentially processed by the host, with AFAs showing greater systemic exposure than BCFAs.

### *Iba* and *Bhy* encode diverse 2-oxoacid:ferredoxin oxidoreductases that act on branched-chain and aromatic substrates

The dominant non-SCFA products formed by *Iba* and *Bhy* were chain-shortened organic acids characteristic of branched-chain and aromatic amino acid oxidation. In anaerobes, such transformations are often initiated by 2-oxoacid:ferredoxin oxidoreductases (OFORs), which convert 2-oxoacids to their corresponding acyl-CoA thioesters while liberating CO_2_ and reducing ferredoxin. Consistent with this, OFORs have established roles in branched-chain amino acid catabolism in *Clostridium sporogenes* (VOR)^28^ and in aromatic amino acid metabolism in diverse anaerobes (e.g., PPFOR/IOR)^29,30^. We therefore asked whether *Iba* and *Bhy* encode OFORs that could account for their product profiles.

Using PSI-BLAST searches seeded with *Moorella thermoacetica* OOR subunits, we identified three candidate OFORs in *Iba* and ten in *Bhy*, with *Bhy* representing the highest OFOR count among strains in our library (**Figure 3A**). Phylogenetic placement using a published OFOR grouping framework^31^ suggested that the *Iba* enzymes distribute across clades consistent with the observed metabolite spectrum, including putative PFOR, VOR, and PPFOR assignments (**Figure 3B**). *Bhy* encoded a larger set including one putative PFOR, one putative OGOR, two putative PPFORs, and multiple putative VORs (**Figure 3B**), suggesting either functional redundancy or specialization across substrates.

**Figure 3.**
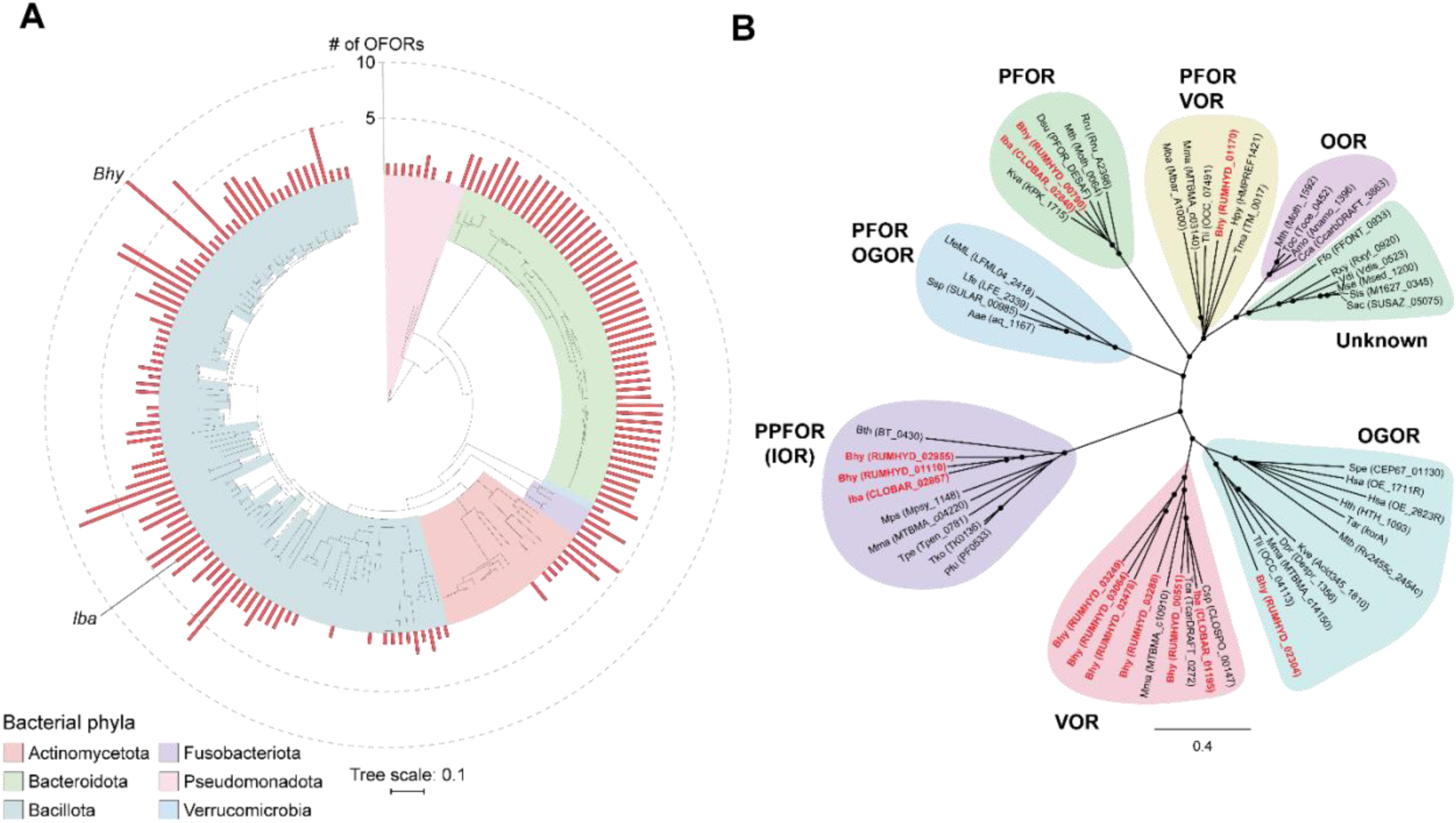
Phylogenetic diversity of gut bacterial OFORs. (A) Distribution of OFORs among bacteria in our strain library with complete or draft genome sequences. The tree is based on RpoB protein sequence alignments, reflecting phylogenetic relationships. Red bars indicate the total numbers of OFORs detected in each strain. (B) Phylogenetic analysis of characterized OFORs and the three OFORs in *Iba* and ten OFORs in *Bhy*. *Iba* and *Bhy* OFORs are indicated in red and bold font.

Because genetic tools are not available for *Iba* or *Bhy*, we heterologously expressed candidate OFOR operons (**Figure S2**; *see methods for cloned genes*) in *E. coli* and measured activity in anaerobic lysates using methylviologen reduction as a readout, coupled to LC-MS verification of select CoA-thioester products (**Figure 4A** and **Figure S3**). As a benchmark, we expressed the genetically validated VOR from *C. sporogenes* (CLOSPO_00146-00149). Reactions containing phenylpyruvate showed a dose-dependent increase in methylviologen absorbance (**Figure S3A**) and produced an LC-MS feature consistent with phenylacetyl-CoA that increased over time (**Figure S3B-C**), confirming that the spectrophotometric signal reflected OFOR-dependent acyl-CoA formation.

**Figure 4.**
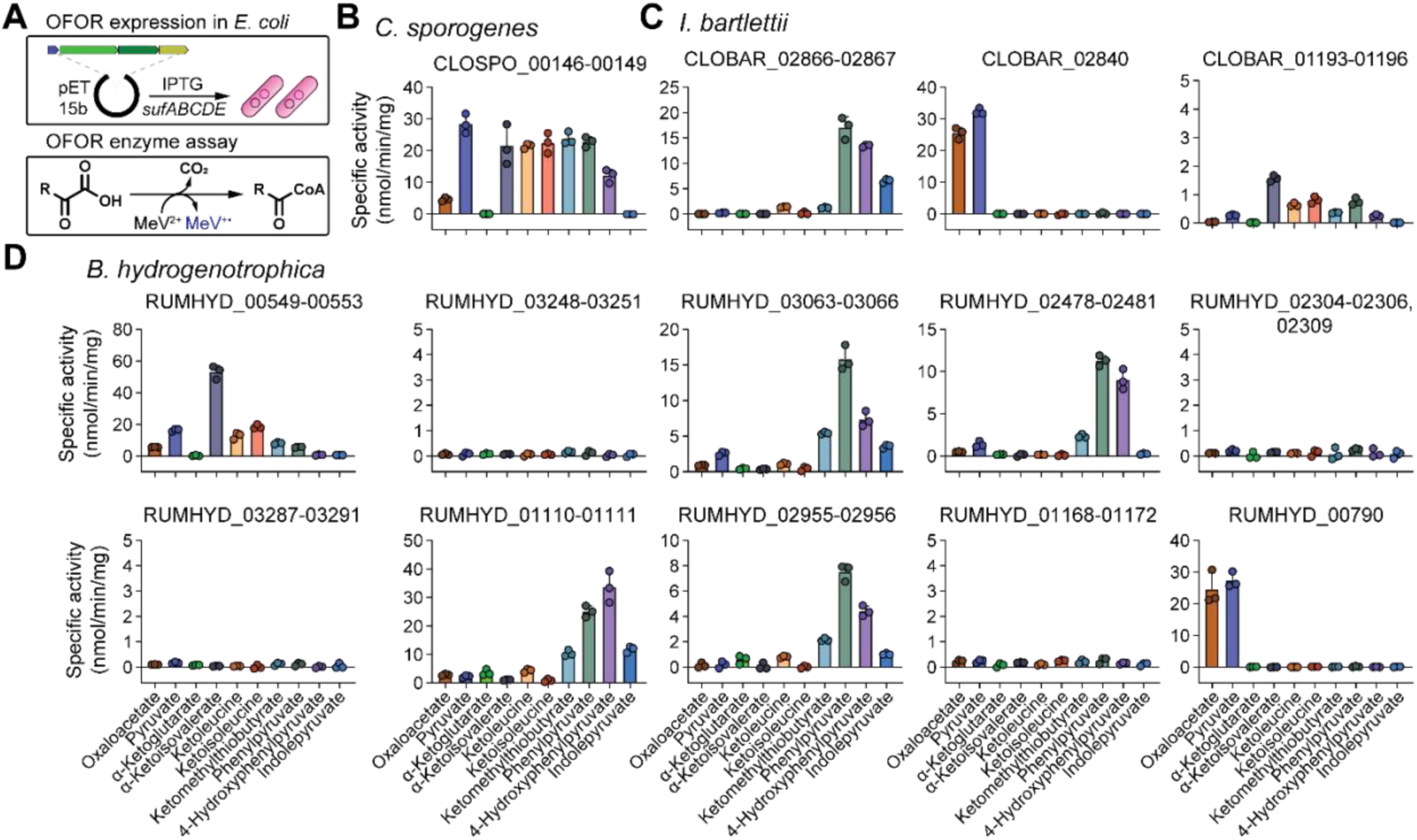
Expression and biochemical analysis of *Iba* and *Bhy* OFORs reveal enzymes with distinct substrate specificities. (A) Schematic of expression and enzyme assays. Genes encoding candidate OFORs were cloned into an IPTG-inducible expression vector and co-expressed in *E. coli* with a vector encoding the iron-sulfur cluster assembly and repair system (*sufABCDE*). Clarified cell lysates were incubated with 2-oxoacids and enzyme activity was assayed anaerobically by reduction of methylviologen, read out spectrophotometrically at an absorbance of 600 nm. (B-D) Substrate specificities for VOR (CLOSPO_00146-00149) from *C. sporogenes* (B), three OFORs from *Iba* (C) and ten OFORs from *Bhy* (D). Genes encoding OFORs are indicated by locus tag IDs. For panels (B)-(D), data are plotted as means + SD for n = 3 replicates with individual values shown.

Using a panel of 2-oxoacid substrates, *C. sporogenes* VOR displayed broad activity, with strong turnover of pyruvate and the branched-chain 2-oxoacids (α-ketoisovalerate, ketoleucine, and ketoisoleucine), substantial activity with the methionine-derived 2-oxoacid ketomethylthiobutyrate and phenylalanine-derived 2-oxoacid phenylpyruvate, and ∼50% relative activity with the tyrosine-derived 2-oxoacid 4-hydroxyphenylpyruvate (**Figure 4B**). While we did not determine kinetic parameters, these activity profiles are consistent with published genetic evidence for a primary role in branched-chain amino acid metabolism and a minor (but significant) activity with phenylalanine^32,28,29^.

When we applied this assay to *Iba* candidates, substrate preferences matched phylogenetic predictions. CLOBAR_02840 preferred pyruvate/oxaloacetate (consistent with a PFOR-like assignment), CLOBAR_01193-01196 favored branched-chain substrates (VOR-like), and CLOBAR_02866-02867 preferred aromatic substrates (PPFOR-like) (**Figure 3B** and **Figure 4C**). Notably, with indolepyruvate, CLOBAR_02866-02867 generated a product consistent with indoleacetyl-CoA (**Figure S3D-E**), implicating this enzyme as a likely entry point from tryptophan-derived 2-oxoacids into an IAA-producing pathway.

The expanded OFOR repertoire in *Bhy* yielded a more complex picture (**Figure 4D**). Several candidates showed no detectable activity in our heterologous system, which may reflect expression/assembly limitations in *E. coli* or substrate preferences outside our test set. Among the six active enzymes, four agreed with phylogenetic assignments: (i) RUMHYD_00790 exhibited PFOR-like specificity toward pyruvate/oxaloacetate; (ii) RUMHYD_02955-02956 and (iii) RUMHYD_01110-01111 both exhibited PPFOR-like specificity toward aromatic substrates; and (iv) RUMHYD_00549-00553 exhibited VOR-like activity toward branched-chain substrates, consistent with prior evidence that branched-chain amino acids induce expression of this locus in *Bhy*^25^. Two enzymes deviated from sequence-based expectations:

RUMHYD_03063-03066 and RUMHYD_02478-02481 are VOR-like by sequence but preferentially acted on aromatic 2-oxoacids. The sequence of the expression vector was confirmed, and protein expression was repeated, yielding similar results. These proteins cluster together on a deeper branch within the VOR-like region of the tree (**Figure 3B**), suggesting a distinct subgroup with aromatic specialization. Unlike the other aromatic-active enzymes, RUMHYD_02478-02481 lacked activity on indolepyruvate.

Together, these data support a model in which *Iba* and *Bhy* use OFORs as gateway enzymes for oxidative metabolism of branched-chain and aromatic amino acids. In this scheme, aminotransferases generate cognate 2-oxoacids, OFORs produce acyl-CoA intermediates, and downstream enzymes convert these to the observed organic acids. Consistent with this pathway architecture, multiple OFOR loci in *Iba* and *Bhy* co-occur with aminotransferases, and homologs of phosphate butyryltransferase, and butyrate kinase (**Figure S2**), a gene context compatible with ATP generation via substrate-level phosphorylation during amino acid oxidation.

*B. hydrogenotrophica* couples amino acid oxidation to acetogenesis via CO_2_ reassimilation.

*Iba* and *Bhy* produce abundant amino acid-derived metabolites consistent with OFOR-mediated oxidative decarboxylation of 2-oxoacids. Because these reactions generate reduced ferredoxin and release CO_2_, they must be paired with an electron-accepting process to maintain redox balance. While many anaerobes couple amino acid oxidation to reduction of a second substrate (e.g., Stickland metabolism), *Iba* and *Bhy* do not accumulate reduced Stickland end products and lack key enzymes for canonical reductive Stickland pathways (e.g., proline/glycine reductases and associated dehydratases).

*Bhy* is a known acetogen capable of autotrophic growth on H_2_/CO_2_^33,34^, and *Iba* encodes a similar complement of WLP genes (**Figure S4**), suggesting that reducing equivalents and CO_2_ released during amino acid oxidation could be recaptured and reduced to acetate. We tested this model using position-specific ^13^C tracing of phenylalanine (**Figure 5A**). During incubation with unlabeled Phe, *Bhy* produced phenylacetic acid (PAA) without detectable labeling. As predicted for oxidative decarboxylation of phenylpyruvate, incubation with 2-^13^C-Phe yielded M+1 PAA, whereas incubation with 1-^13^C-Phe yielded predominantly unlabeled (M+0) PAA (**Figure 5B**), consistent with loss of the C1 label as CO_2_. Notably, acetate M+1 and M+2 isotopologues increased specifically in the 1-^13^C-Phe condition relative to unlabeled Phe and 2-^13^C-Phe (**Figure 5C-D**), indicating incorporation of the decarboxylated carbon into acetate.

**Figure 5.**
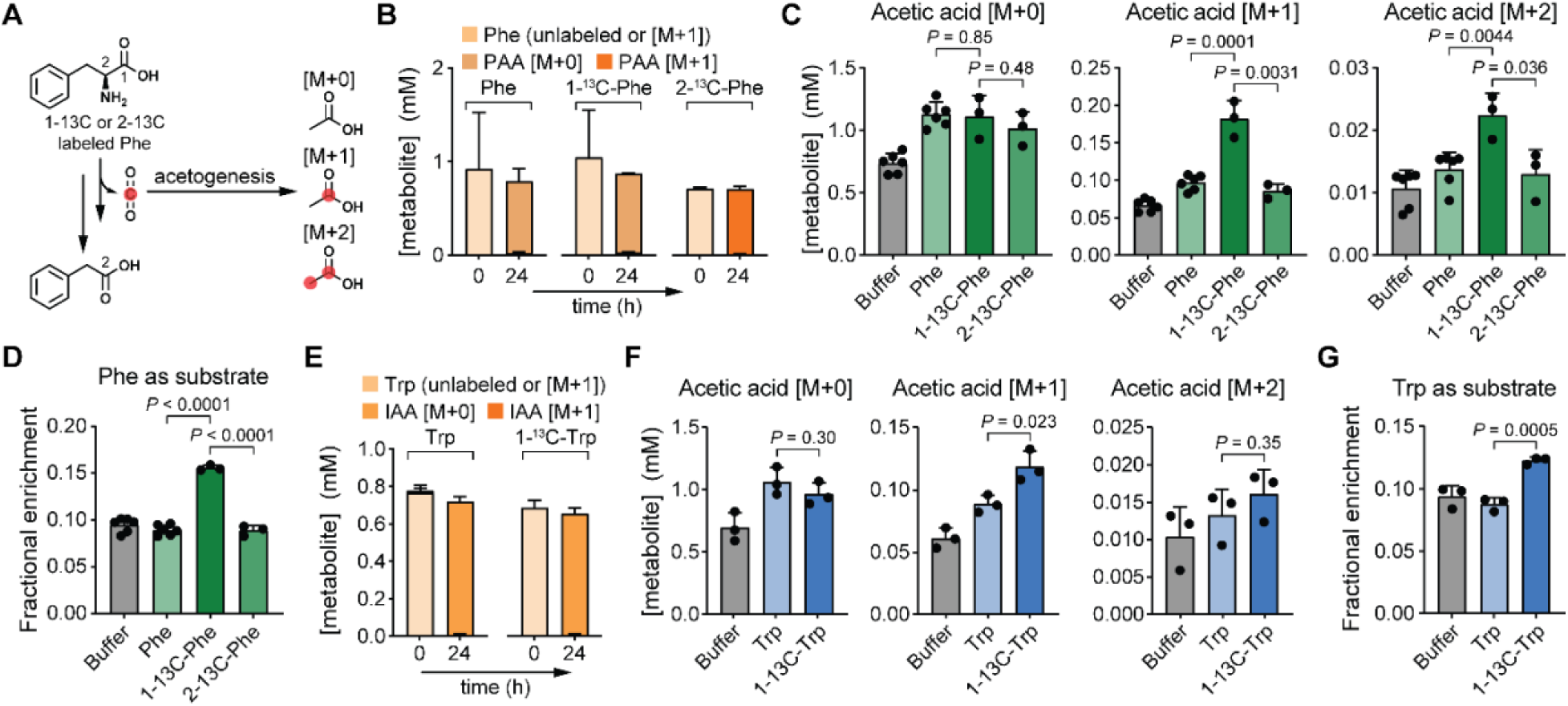
CO_2_ liberated from Phe and Trp are reassimilated into acetate by the acetogen, *Bhy*. (A) Schematic of position-specific Phe isotope tracing in *Bhy*. Cell suspensions were incubated with ^13^C-Phe labeled at either the 1- or 2-carbon and acetate isotopes were quantified by LC-MS after 24 hours. (B) Unlabeled or labeled Phe consumption by *Bhy* quantified by LC-MS over time. PAA isotopes are represented as stacked bar plots. (C) Acetate isotopes quantified by LC-MS at 24 hours following incubation of *Bhy* cell suspensions with either buffer, unlabeled Phe, 1-^13^C-labeled Phe, or 2-^13^C-labeled Phe. (D) Fractional enrichment of M+1 and M+2 acetate isotopes under conditions in (C). (E) Unlabeled or labeled Trp consumption by *Bhy* quantified by LC-MS over time. IAA isotopes are represented as stacked bar plots. (F) Acetate isotopes quantified by LC-MS at 24 hours following incubation of *Bhy* cell suspensions with either buffer, unlabeled Trp, or 1-^13^C-labeled Trp. Note, 2-^13^C-labled Trp was not commercially available. (G) Fractional enrichment of M+1 and M+2 acetate isotopes under conditions in (F). For panels (D) and (G), fractional enrichment was calculated as ([M+1] + [M+2]) / ([M+0] + [M+1] + [M+2]). Note that some [M+2] acetate isotope values dropped below the calibration curve range and are less accurate (see **Table S10**). For panels (B)-(G), data are plotted as means + SD from n = 3 replicates and *P*-values are from an unpaired student’s t-test.

We observed a similar pattern with tryptophan: conversion to indoleacetic acid (IAA) proceeded without label retention in IAA from 1-^13^C-Trp (**Figure 5E**), while acetate became enriched primarily as M+1 (**Figure 5F-G**). Together, these results support a model in which *Bhy* reassimilates CO_2_ released during amino acid oxidation into acetate. The reducing equivalents released as ferredoxin via OFOR activity may also enter the WLP to enable redox balancing during amino acid catabolism. Direct genetic tests (e.g., WLP disruption) would further establish this coupling, but targeted mutagenesis is not currently feasible in *Bhy*.

## Discussion

Microbial fermentation of carbohydrates and amino acids in the gastrointestinal tract generates dozens of small molecules that act locally in the gut and, in some cases, enter circulation to affect distal organs. Linking these metabolites to the specific microbes and pathways that produce them is essential for understanding how microbiome composition shapes host physiology and for designing rational dietary or microbiota-targeted interventions. Here we focused on indole-3-acetic acid (IAA), a tryptophan-derived microbial metabolite with reported immunomodulatory effects via AhR-dependent and AhR-independent mechanisms.

Our cultivation-based analysis across a large library of 206 human gut bacterial isolates shows that robust IAA production is an uncommon phenotype that is restricted to a few bacteria within the Bacillota phylum. These findings are consistent with a previous study which evaluated 26 gut bacterial strains and identified only a single robust IAA producer (*Intestinibacter bartlettii*)^23^, which also emerged as a strong producer in our screen. Given that IAA is frequently detected in human stool, plasma, and urine^35,36,37,38^, our data imply that IAA abundance *in vivo* may be disproportionately controlled by a small number of taxa rather than a functionally redundant set of organisms. One consequence is vulnerability: ecological pressures that deplete these strains (e.g., antibiotics, intestinal inflammation, etc.) could cause drops in IAA levels, consistent with observations of reduced stool IAA in inflammatory bowel disease^12^.

Conversely, the smaller producer pool also suggests an actionable therapeutic angle: restoring IAA production may require only a finite set of high-capacity strains. Notably, *B. hydrogenotrophica* (developed as MRx1234/Blautix^TM^) is being evaluated clinically for irritable bowel syndrome^39^ and IAA production (alongside other metabolic outputs) could plausibly contribute to its mechanism.

A second key finding is that, even in these “high-IAA” strains, IAA was not the dominant metabolic output. Instead, *Iba* and *Bhy* accumulated a broad suite of branched-chain and aromatic organic acids at concentrations that far exceeded IAA. This indicates that tryptophan-to-IAA conversion is best viewed as one branch within a broader amino acid-oxidizing program. *In vitro*, supplying carbohydrates suppressed amino acid oxidation and its downstream products, consistent with carbon catabolite control. If similar regulation operates *in vivo*, then carbohydrate availability (whether delivered directly by diet or indirectly via cross-feeding) could shift flux away from amino acid-derived products (including IAA) and toward carbohydrate fermentation end products. Testing this hypothesis will require *in vivo* studies that explicitly vary fermentable carbohydrate and protein availability while measuring metabolite outputs and strain abundance.

Gnotobiotic colonization with *Iba* further showed that microbial products differ markedly in their capacity to reach systemic circulation and in how extensively they are remodeled by host metabolism. BCFAs were abundant in intestinal contents yet were largely extracted between portal and peripheral plasma, whereas aromatic fatty acids (AFAs) and especially their glycine conjugates increased in plasma and urine, consistent with appreciable systemic exposure and host processing. These results emphasize that the impact of microbial metabolism on the host depends on two separable determinants: (i) microbial biosynthetic capacity and its regulation by diet and ecology, and (ii) host absorption, hepatic clearance, and conjugation/excretion pathways that shape which microbial metabolites persist in circulation. The near-complete hepatic extraction of BCFAs suggests that these compounds may contribute to host metabolism (e.g., via hepatic BCFA catabolic pathways), whereas the appearance of aromatic acylglycines in urine supports the use of these conjugates (and, in humans, the corresponding glutamine conjugates) as practical, noninvasive readouts of gut microbial aromatic metabolism.

Our data identify 2-oxoacid:ferredoxin oxidoreductase (OFOR) enzymes as a central gateway connecting amino acid availability to production of IAA and chemically related metabolites. Phylogenetic and biochemical characterization of candidate OFOR operons from *Iba* and *Bhy* demonstrated activities spanning pyruvate/oxaloacetate (PFOR-like), branched-chain 2-oxoacids (VOR-like), and aromatic 2-oxoacids (PPFOR-like), consistent with the diverse organic acids observed in culture. Importantly, we observed aromatic substrate preferences not only in PPFOR-like enzymes but also in VOR-like loci in *Bhy*, suggesting that sequence-based annotations may underpredict aromatic catabolic capacity in some gut anaerobes. Despite targeted comparisons among OFORs, we did not identify clear sequence or structural motifs that reliably predicted indolepyruvate oxidation (data not shown), underscoring the need for broader structure-function mapping across this diverse enzyme family.

Extensive OFOR-dependent amino acid oxidation also raises a physiological question: how do these organisms conserve energy and maintain redox balance when amino acids become major substrates? Two, non-mutually exclusive mechanisms likely contribute. First, many OFOR loci in *Iba* and *Bhy* co-occur with homologs of phosphate butyryltransferase and butyrate kinase (**Figure S2**), consistent with ATP generation via substrate-level phosphorylation downstream of acyl-CoA formation. Second, our position-specific ^13^C tracing indicates coupling between oxidative amino acid metabolism and reductive acetogenesis, supporting an “intracellular syntrophy” model in which reducing equivalents generated during oxidation are consumed by the Wood-Ljungdahl pathway to support redox balance and energy conservation^40^. This framework expands the ecological view of gut acetogens beyond obligate dependence on external syntrophic partners, positioning them as capable of coupling oxidation and CO_2_ reduction within a single organism to remain metabolically self-sustaining.

Together, these findings suggest concrete directions for modulating gut microbial metabolite output. *In vitro*, fermentable carbohydrate supplementation suppressed amino acid oxidation and its downstream organic acid products in *I. bartlettii* and *B. hydrogenotrophica*, consistent with strong nutrient-dependent regulation of this metabolic program. However, translating these relationships to the gut ecosystem is not straightforward. Fermentable carbohydrates can reshape community structure and substrate partitioning, potentially redirecting tryptophan and other amino acids between taxa and end products via cross-feeding interactions. For example, shifts in indole versus indole-3-propionic acid producing pathways have been observed in defined systems^41^. Accordingly, carbohydrate availability likely serves as one of several variables, along with protein form and accessibility, intestinal transit, and community context, that may tune OFOR-linked amino acid oxidation and IAA output *in vivo*.

At the community level, the rarity of high-capacity IAA producers implies that relatively small changes in strain abundance could drive large shifts in IAA and related metabolites, motivating strain-resolved monitoring and targeted interventions. Finally,

OFOR and WLP nodes represent rational control points for future work, whether as diagnostic markers of pathway activity, targets for small-molecule inhibition, or design principles for engineered consortia. Key limitations remain: our enzyme activity measurements rely on heterologous expression and lack kinetic parameters, and direct genetic tests of pathway coupling in *Iba* and *Bhy* are not yet feasible. Addressing these gaps, together with *in vivo* studies that manipulate diet and community context, will be essential for translating these mechanistic insights into predictable strategies to tune microbiota-derived metabolism in the host.

## Methods

### Bacterial strains and culture conditions

#### Strains and general culture conditions

The bacterial strains used in this study are listed in **Table S1**. Glycerol stocks (20%) were prepared anaerobically, and strains were sealed and stored at -80 °C. All bacteria were cultured at 37°C using a Coy type B anaerobic chamber containing a gas mix with the composition of: 5% hydrogen, 10% carbon dioxide, and 85% nitrogen. Hydrogen levels were maintained at 3.3% using an anaerobic gas infuser. All media and plasticware were pre-reduced in the anaerobic chamber for at least 24 hours before use. *E. coli* strains for protein overexpression and purification were cultured aerobically using LB plates and broth, with temperature and antibiotic selection varying depending on the protocol.

#### Screening of bacterial strain library

Bacterial strain library screening was performed anaerobically in 2 mL 96-deep well plates. Glycerol stocks were first inoculated into rich media and incubated at 37 °C for 24 h. Cultures were subsequently diluted 1:10 into fresh medium supplemented with 1 mM tryptophan and incubated for an additional 48 h. Supernatants were harvested by centrifugation at 5,000 x *g* for 25 min at 4 °C. Aliquots of the supernatants were then transferred to 96-well microtiter plates, sealed tightly, and stored at -80 °C until sample preparation for liquid chromatography-mass spectrometry (LC-MS) analysis.

#### Metabolic profiling and growth measurement of Iba and Bhy in multiple media

For metabolic profiling on multiple media, *Iba* and *Bhy* were first streaked on RCM plates and incubated at 37 °C for 48 hours. Single colonies were inoculated into Chopped Meat Medium (ChopM; Anaerobe Systems, AS-8115) and grown for 24 hours. Saturated cultures were subcultured into fresh ChopM media at a 25-fold dilution and incubated overnight (to increase volume of cell culture). These overnight cultures were centrifuged anaerobically at 7,000 x *g* for 10 min and gently rinsed with buffer twice, making sure to fully resuspend the cell pellets. Cells were resuspended in buffer to the original volume and diluted 20-fold into respective media. Aliquots were taken across the time points and split between two 96-well plates, one for OD_600_ measurement via a plate reader (BioTek Synergy H1) and the other for harvesting supernatant after centrifugation at 5,000 x *g* for 10 min. Supernatants were stored at -80 °C until LC-MS analysis.

#### Concentrated cell assay for stable isotope tracing

*Bhy* DSM 10507 was streaked on an RCM plate and incubated at 37 °C for 48 hours. Single colonies were inoculated in rich media for 24 hours, subcultured in fresh rich media in a 50-fold dilution, then incubated overnight. Cultures were centrifuged anaerobically at 7,000 x *g* for 10 min and gently rinsed with buffer twice. Cells (final OD_600_ = 2) were incubated with 1 mM of substrate in a sealed LC-MS vial to reduce environmental CO_2_ exchange. Substrates were aliquoted into open LCMS vials, and the screw top was closed immediately after adding the concentrated cells. After 24 hours, samples were transferred to pre-chilled 1.5 mL Eppendorf tubes and centrifuged at >20,000 x *g* for 1.5 min. Supernatants were collected and stored at -80 °C until LC-MS analysis.

## Animal Husbandry

All mouse experiments were conducted under a protocol approved by the Stanford University Institutional Animal Care and Use Committee. Experiments were performed on male and female gnotobiotic Swiss Webster germ-free mice (aged 11-15 weeks, n = 8; 5 male, 3 female) originally obtained from Taconic Biosciences (*Mus musculus*, Tac:SW) and maintained in aseptic flexible film isolators. Mice were fed a standard chow (LabDiet, catalog no. 5K67) and sterile water with access to food and water *ad libitum* in a facility on a 12-hour light/dark cycle with temperature controlled between 20-22 °C and humidity between 40-60%.

*Mono-colonization of gnotobiotic mice with* Iba. Before the experiment, the sterility of the mice was confirmed by incubating a fecal pellet in rich media in both aerobic and anaerobic conditions. *Intestinibacter bartlettii* was cultured overnight anaerobically at 37 °C in Mega media and administered to mice via intragastric gavage. Fresh fecal pellets were obtained from individual mice, and fresh urine was collected from individual mice into sterile tubes. A facial vein puncture was used to collect samples of circulating blood (∼100uL) into a tube containing the anticoagulant, EDTA (final concentration ∼15 mM). Blood samples were centrifuged at 1,500 x *g* for 10 min at 10°C, and the plasma was transferred to a new tube. Cecal contents were collected post-euthanasia via CO_2_ asphyxiation and cervical dislocation. All samples were stored at -80 °C.

*Portal vein sampling.* Mice were anaesthetized with a solution of ketamine and xylazine using an IP injection in the lower left abdomen, for a final concentration of 100 mg/kg ketamine and 10 mg/kg xylazine per mouse. Breathing rate and the color of mucous membranes were constantly monitored for signs of distress. Responsiveness was determined by the toe pinch test, and after the animal was determined to be in a deep anesthetic plane, an abdominal incision was made. Organs were displaced to identify the portal vein, which was catheterized, and ∼100 μL of blood was collected into a tube containing the anticoagulant, EDTA (final concentration ∼15 mM). Immediately after sample collection, mice were humanely euthanized by cervical dislocation. Samples were centrifuged at 1,500 x *g* for 10 min at 10 °C, and the plasma was transferred to a new tube. Samples were stored at -80 °C.

### Protein overexpression and purification

#### Cloning OFORs into the overexpression plasmid, pET-15b

Enzymes were heterologously expressed using the pET-15b plasmid, which contains an IPTG-inducible promoter and the β-lactamase gene which confers carbenicillin resistance. Gene sequences were first codon optimized for *E. coli* to increase the chance of successful expression, and ribosome binding sites were reverted to the original sequence, if necessary, to maintain functionality. All subunits in a multi-subunit PFOR enzyme were overexpressed in the same vector, in the same layout as in the genome. The empty vector control contains the pET-15b plasmid without any insertions. These sequences were then synthesized as gBlocks by IDT (**Table S2**). The pET-15b plasmid backbone was linearized using BamHI-HF, dephosphorylated with shrimp alkaline phosphatase (rSAP), and gel purified. Gibson assembly was used to assemble the pET-15b backbone with the gBlocks using NEBuilder HiFi DNA Assembly Master Mix. These Gibson assemblies were transformed into *E. coli* Stbl4 by electroporation and grown on LB-carbenicillin plates (100 μg/mL) to select for transformants. Plasmids were extracted from the transformants and sent for whole-plasmid sequencing.

#### Heterologous expression in E. coli

Because OFORs contain iron-sulfur clusters, we chose to co-express these genes with the *E. coli* iron-sulfur cluster assembly system *(sufABCDE*), encoded within the chloramphenicol resistant plasmid pPH151. OFORs cloned within pET-15b were co-transformed with pPH151 into *E. coli* BL21 (DE3) by electroporation and were selected on LB plates supplemented with carbenicillin (100 μg/mL) and chloramphenicol (25 μg/mL). Colonies were inoculated into ∼11 mL of LB broth supplemented with the same antibiotics and incubated at 37 °C aerobically with shaking overnight. Approximately 10 mL of overnight cultures were then inoculated in 1L of LB + antibiotics + FeCl_3_ (200 μM) in 2.8L baffled Fernbach flasks and grown at 37 °C with vigorous aeration on an orbital shaker (225 rpm) until an OD ∼0.4. Cultures were then cooled on wet ice to ∼16 °C, at which point IPTG (final concentration 0.1 mM) and L-cysteine (0.6 mM) were added, and cultures were incubated at 16 °C overnight with vigorous aeration. Cells were harvested by centrifugation at 5,000 x *g* for 15 min at 4 °C and cell pellets were stored at -80 °C until cell lysis.

#### Cell lysis and preparation of cell extracts

All cell lysis steps for OFORs were performed in an anaerobic chamber. Buffers were deoxygenated by heating until boiling, then sonicated in a sonicating bath for 30 minutes, and then β-mercaptoethanol (5 mM final) was added as a reducing agent and buffers were brought into the chamber ready for use. Cell pellets were resuspended in a binding buffer containing sodium phosphate (50 mM) and sodium chloride (300 mM) and lysed via sonication while suspended in an ice bath. Cell lysate was centrifuged at 7,197 x *g* for 30 min at 4 °C, and supernatants were transferred to a new tube. Total protein concentrations in OFOR clarified cell lysates were measured using Bio-Rad DC Protein Assay Kit II, then lysates were combined with pre-reduced 80% glycerol (20% final) and aliquoted into LC-MS vials with a glass insert, sealed tightly to reduce exposure to oxygen, and stored at -80 °C.

## Enzyme assays

All assays were performed anaerobically with pre-reduced substrates, buffers, reagents, and plasticware. Hermetically sealed cell lysates were kept on ice and brought into the anaerobic chamber immediately before use.

### Testing OFOR substrate specificity using methyl viologen

Cell lysates from *E. coli* overexpressing OFOR enzymes were incubated in a potassium phosphate buffer (100 mM) with thiamine pyrophosphate (0.4 mM), coenzyme A (0.1 mM), MgCl_2_ (2.5 mM), methyl viologen (1 mM), and substrate (5 mM) at 37 °C in a plate reader (BioTek Synergy H1) monitoring absorbance at 600 nm (with 1 cm pathlength correction) every 51 seconds for 1 hour. Multiple dilutions of each cell lysate were tested to identify the dilution that resulted in activity in a linear range. This dilution was then used to test activity for various substrates in triplicate. Enzyme specific activity was calculated using the Beer-Lambert Law. In short, the slope of the linear range of the reaction progress curve was divided by the extinction coefficient of methyl viologen (13,700 cm^-1^ M^-1^), adjusted by the appropriate dilution factor, and divided by the total protein concentration of the cell lysate.

#### Sample preparation for the detection of amino acids, acetic acids, and pathway intermediates by liquid chromatography-mass spectrometry (LC-MS)

### Extraction of metabolites from liquid samples

Supernatants from in vitro experiments, as well as plasma and urine samples, were mixed with internal standards in 96-well plates before being mixed with an acetonitrile/methanol (3:1) solution for extraction and protein precipitation. Plates were covered with a Costar 96-well grid lid to prevent evaporation and centrifuged at 5,000 x *g* for 20 min. Supernatants were transferred to a new plate and either diluted for LC-MS analysis or used for derivatization. Supernatants collected from enzyme assays were directly diluted and run on the LC-MS.

### Extraction of metabolites from fecal/cecal samples

Fecal and cecal samples were weighed out into screw-top microcentrifuge tubes with an equivalent mass of glass beads. Internal standards and acetonitrile/methanol solution (3:1) were added to the tubes, and samples were homogenized with a bead beater at a rate of 25/s for 30 min at 4 °C. Tubes were centrifuged at 13,000 x *g* for 5 min at 4 °C, and supernatants were transferred to a 96-well plate before undergoing derivatization.

### 3-Nitrophenylhydrazine (3-NPH) derivatization

3-NPH derivatization is helpful for measuring molecules that contain a free carboxylic acid, as it enables chemically diverse compounds to be retained on C18 chromatographic media and boosts signal during electrospray ionization. Extracted supernatants were mixed (1:1) with a solution containing 100 mM 3-NPH, 60 mM N-(3-dimethylaminopropyl)-N’-ethylcarbodiimide (EDC), and 3% pyridine in 25% acetonitrile (diluted in water). Samples were incubated in a thermomixer at 600 rpm for 60 min at 40 °C and then quenched in a solution containing 10% acetonitrile (diluted in water) and 0.02% formic acid.

### Dansyl chloride (DNS) derivatization

DNS derivatization enhances compound retention on C18 columns and boosts signal for molecules that contain primary or secondary amines. Extracted supernatants were mixed (1:15) with a solution containing 50 mM DNS (in 50% acetonitrile) and 100 mM sodium carbonate/bicarbonate (pH 9.8). Sample were incubated in a thermomixer at 300 rpm for 60 min at 25 °C, then 10% ammonium hydroxide (1:10 total volume) was added and incubation was continued for an additional 5 min to consume extra DNS. Samples were briefly centrifuged at 1,000 x *g* for 1 min at room temperature and then mixed (1:25) with a solution of 40% acetonitrile (diluted in water) and 0.01% formic acid.

### Liquid chromatography-mass spectrometry (LC-MS)

Quantification of metabolites by LC-MS During this study, four different LC-MS conditions were used: C18 positive/negative underivatized (for urinary creatinine and arylpyruvates), Poroshell positive underivatized (for CoA species), 3-nitrophenylhydrazine (NPH) derivatized C18 negative (for fatty acids), and dansyl chloride (DNS) derivatized C18 positive (for select amino acids). An overview of the general method is provided here and the specific instrument parameters for the different analytical methods are provided in **Table S3**. Samples were injected via a refrigerated autosampler into mobile phase, chromatographically separated by an Agilent 1290 Infinity II UPLC system and detected using an Agilent 6545XT Q-TOF system equipped with a dual jet stream electrospray ionization source operating under extended dynamic range (1,700 m/z). MS1 spectra were collected in centroid mode, and peak assignments in samples were made on the basis of comparisons of retention times and accurate masses from authentic standards (when available) using MassHunter Quantitative Analysis v.10.0 software from Agilent Technologies. Compounds were quantified from calibration curves constructed with authentic standards (when available) using isotope-dilution mass spectrometry with appropriate internal standards (**Table S4**).

#### Characterization of phenylacetyl-CoA and indoleacetyl-CoA by MS/MS

Samples containing LC-MS features consistent with phenylacetyl-CoA and indoleacetyl-CoA were subjected to targeted MS/MS. Targeted methods (Poroshell positive mode) were run with predefined precursor m/z for phenylacetyl-CoA ([M+H]+ 886.1649, RT = 3.09 min) or indoleacetyl-CoA ([M+H]+ 925.1759, RT = 3.16 min) with retention time window of 0.3 min, narrow isolation width, fixed collision energy of 20 eV and acquisition time of 400 ms per spectrum. Spectra were manually inspected using Agilent MassHunter Qualitative Analysis (v.10.0.10305.0) and the cleanest spectra centered closest to the maximum peak area for the precursor compound were exported as .txt files and normalized to the maximum fragment ion intensity.

### Bioinformatics analysis

#### Identification of OFORs in our strain library

We performed local BLASTp searches (Geneious Prime v2025.1.2) of our strain library using either the using the alpha and beta subunits of *Moorella thermoacetica* OOR (Locus tag IDs, Moth_1592 and Moth_1591^42^) or the *Pyrococcus furiosus* IOR (Locus tag ID PF0533^30^). Hits from these two BLASTp searches were retrieved, de-replicated, and the number of hits for each bacterial strain in our library were counted. To generate the phylogenetic tree, RpoB protein sequences were retrieved from 187 organisms in our library, uploaded into Geneious Prime, and a Muscle alignment was performed with default parameters. Gaps were then manually trimmed, and sequences were re-aligned with Muscle. A neighbor joining unrooted phylogenetic tree was generated from the alignment and exported as a newick file, then uploaded into the interactive tree of life (iTOL) where OFOR hits were displayed as a bar chart on the outer ring of the tree. NCBI accession number for OFOR hits are provided in **Table S5**.

#### Phylogenetic analysis of OFORs

To generate a phylogenetic tree for the OFOR protein family, we followed a previously published approach^31^, with some minor modifications. First, a subset of sequences for OFOR alpha and beta subunits analyzed in Gibson et al. were retrieved from GenBank. Next, we searched for OFOR homologs in the genomes of *I. bartlettii* (NCBI GenBank assembly GCA_000154445.1) and *B. hydrogenotrophica* (NCBI GenBank assembly GCA_000157975.1) using the alpha and beta subunits of *Moorella thermoacetica* OOR (Locus tag IDs, Moth_1592 and Moth_1591^42^). Routine BLAST searches failed to return all possible homologs, so we used position-specific iterative BLAST (PSI-BLAST) to rebuild the similarity matrix and perform a second round of BLAST. This identified three putative OFORs in *I. bartlettii* and ten putative OFORs in *B. hydrogenotrophica*. Due to the incomplete nature of the *B. hydrogenotrophica* genome assembly, we noticed several partial sequences for OFOR subunits. To enable phylogenetic analysis, we manually curated these proteins as follows: (i) *B. hydrogenotrophica* OFORs were used as queries in a BLASTp search of GenBank and homologs with 100% amino acid identity from other genome assemblies were identified, (ii) Amino acid sequence alignments were performed and *B. hydrogenotrophica* OFOR subunits that showed reduced sequence coverage compared to their homologs were identified. (iii) We then substituted full length homologs from other genome assemblies into the list of sequences to be used for phylogenetic analyses.

Initial multiple sequence alignments were performed with MUSCLE using the PPP algorithm as implemented in Geneious Prime (v2025.1.2). Based on these initial alignments, we concatenated the alpha and beta subunits for genes colocalized on the chromosome into single proteins for each homolog. We then performed another MUSCLE alignment, manually trimmed sequence gaps, and generated an unrooted neighbor-joining phylogenetic tree with Geneious Prime using the Jukes-Cantor distance model and bootstrap resampling with 100 replicates. The tree was exported as a .pdf file and edited in Adobe Illustrator (v29.5). All concatenated OFOR sequences used to generate the alignment and tree are provided in **Table S6**.

#### Gene cluster searches using MultiGeneBlast

We used the tool, MultiGeneBlast, to look for homologs of the *porA* genes from *C. sporogenes* (CLOSPO_00146-00149) and the Wood-Ljungdahl Pathway genes from *C. ljungdahlii* (CLJU_c37520-37670) in the genomes of *Iba* (Genome Assembly ASM15444v1) and *Bhy* (Genome assembly ASM15797v1). Gene clusters were exported to Adobe Illustrator for visualization.

## Data availability

All targeted metabolomics data are provided in **Tables S7-S10**. Strains and plasmids generated in this study are available from the corresponding author without restriction.

## Acknowledgements

We thank Arianna Burke, Alejandra Dimas, and Marissa Jasper for help with gnotobiotic experiments. We thank Tyler Grove for sharing the pPH151 plasmid. This work was funded in part by National Institutes of Health (NIH) grants K08-DK110335 (D.D.), R35-GM142873 (D.D.), R01-AT011396 (D.D.), the Stanford Microbiome Therapies Initiative (D.D.), an OHF-ASN Foundation for Kidney Research Career Development Award (D.D.), a Stanford Medicine Dean’s Postdoctoral Fellowship (Z.Z.), a Stanford Medicine Children’s Health Center for IBD and Celiac Disease Postdoctoral and Early Career Support Award (Z.Z.), and the Stanford Microbiology and Immunology Department NIH T32-AI007328 (M.E.D).

## Author contributions

M.E.D., Y.L., and D.D. conceived and designed the project; Y.L. performed strain library cultures in Trp-supplemented media; M.E.D. performed all *in vitro* experiments with *Iba* and *Bhy* and performed sample preparation for LC-MS and analyte quantitation; D.D., and Z.Z. performed LC-MS method development and acquired LC-MS data; M.E.D. performed gnotobiotic experiments with help from S.K.H.; D.D. performed phylogenetic analyses; M.E.D. cloned enzymes, expressed proteins and performed anaerobic assays with help from D.D.; M.E.D. and D.D. prepared figures and wrote the manuscript with input from all co-authors.

## Competing interests

The authors declare no competing interests.

---

**Figure S1.**
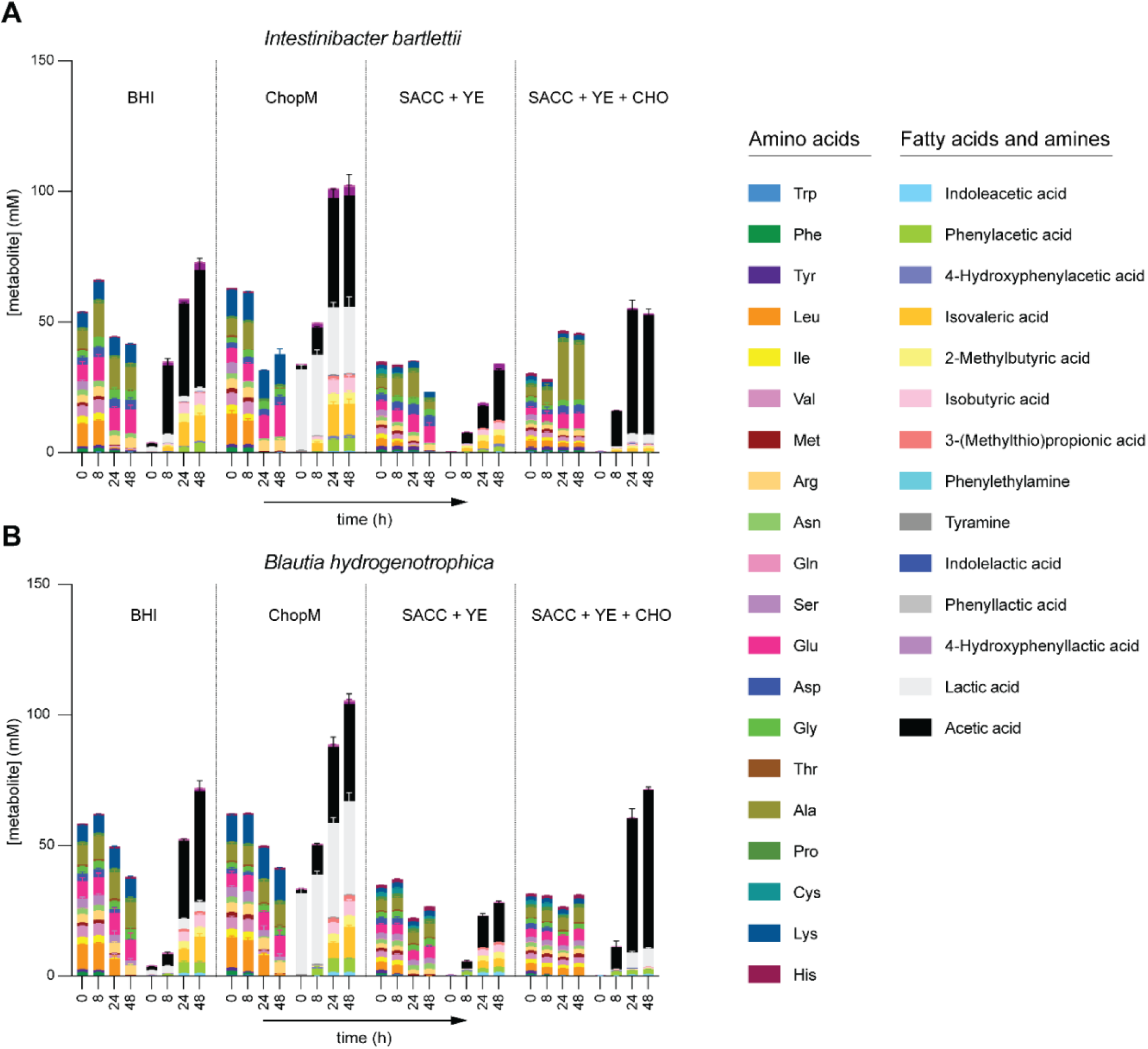
Comprehensive metabolic profiling of *Iba* and *Bhy* in four different media. (A-B) LC-MS time-course profiling of *Iba* (A) and *Bhy* (B) grown in four media, showing amino acid consumption and fatty acid production over time. Data are plotted as means + SD from n = 3 replicates.

**Figure S2.**
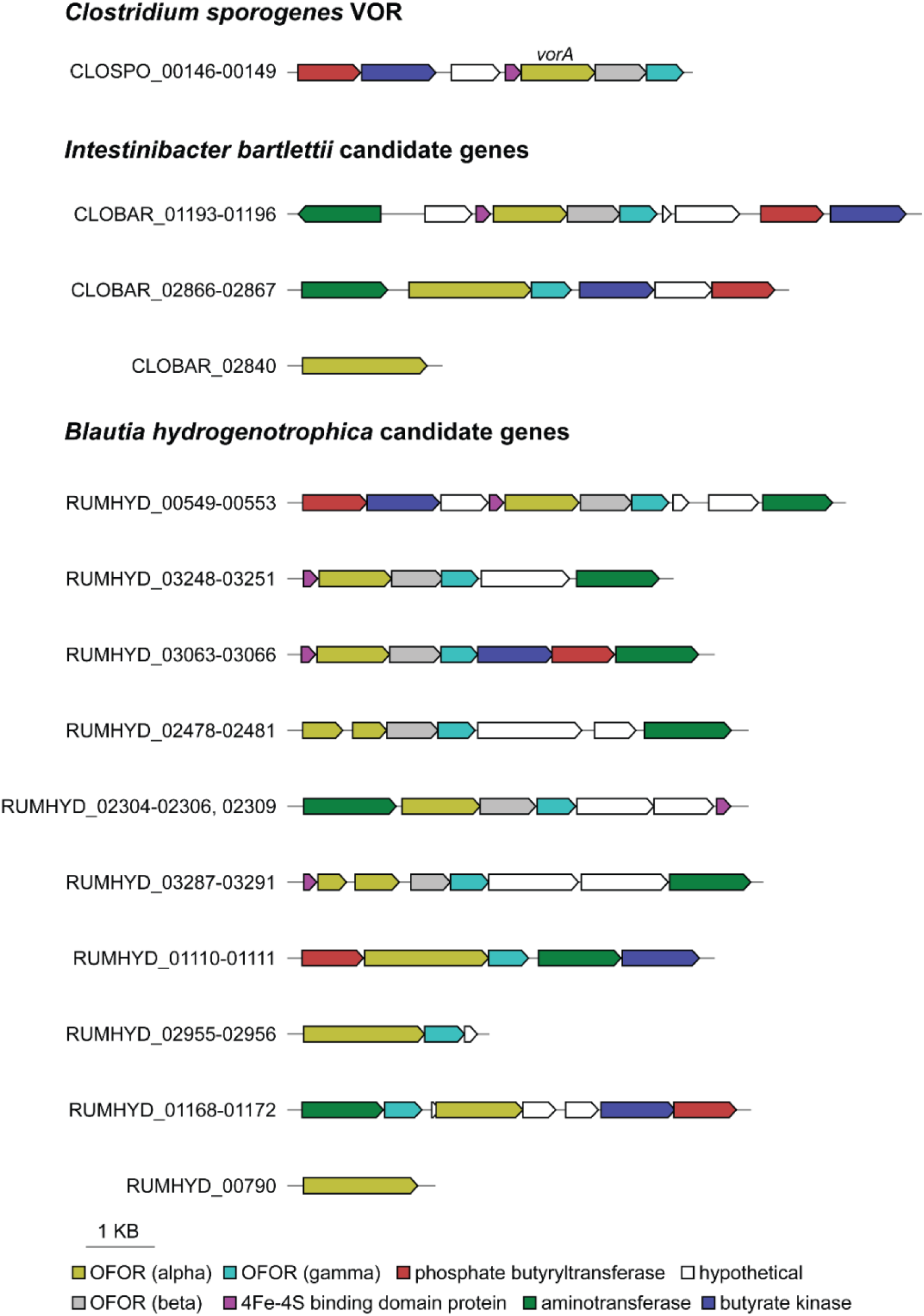
Gene clusters for the known VOR from *C. sporogenes* and candidate OFORs from *Iba* and *Bhy*. The *C. sporogenes* VOR gene cluster with linked genes was used as a query to search the genome sequences of *Iba* and *Bhy* using MultiGeneBlast with an amino acid percent identity cutoff of 15%. The OFOR homologs (CLOBAR_02840 and RUMHYD_00790) were not identified as hits from *C. sporogenes* VOR and they were manually retrieved from the *Iba* and *Bhy* genomes.

**Figure S3.**
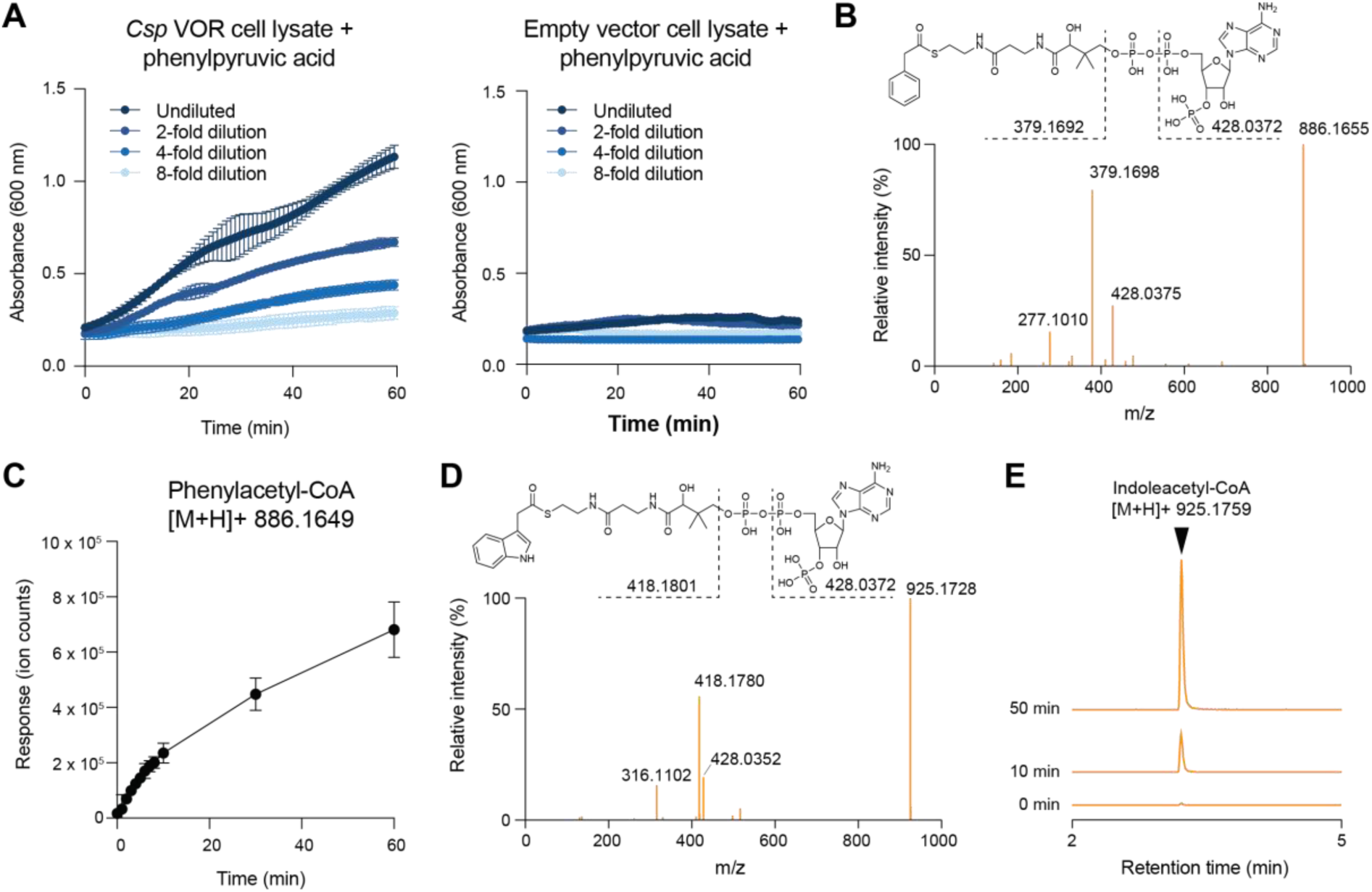
Supporting data for enzyme assays and identification of phenylacetyl-CoA and indoleacetyl-CoA. (A) Methyl viologen reduction progress curves for serial dilutions of *E. coli* cell extracts expressing the *C. sporogenes* VOR or empty vector control, incubated with phenylpyruvic acid. (B) MS/MS spectrum of a candidate phenylacetyl-CoA feature detected in *E. coli* extracts expressing *C. sporogenes* VOR after incubation with phenylpyruvic acid ([M+H]+ 886.1649, RT = 3.09 min). (C) Time-course accumulation of phenylacetyl-CoA in *E. coli* extracts expressing *C. sporogenes* VOR, incubated with phenylpyruvic acid. (D) MS/MS spectrum of a candidate indoleacetyl-CoA feature detected in *E. coli* extracts expressing *Iba* CLOBAR_02866–02867 after incubation with indolepyruvate ([M+H]+ 925.1759, RT = 3.16 min). (E) Extracted ion chromatograms showing an increase in the indoleacetyl-CoA feature ([M+H]+ 925.1759, RT = 3.16 min) over time during incubation of *E. coli* extracts expressing *Iba* CLOBAR_02866–02867 with indolepyruvic acid. For (B) and (D), diagnostic acyl-CoA fragmentation ions are annotated; observed fragment m/z values agree with the predicted acyl-CoA cleavage pattern.

**Figure S4.**
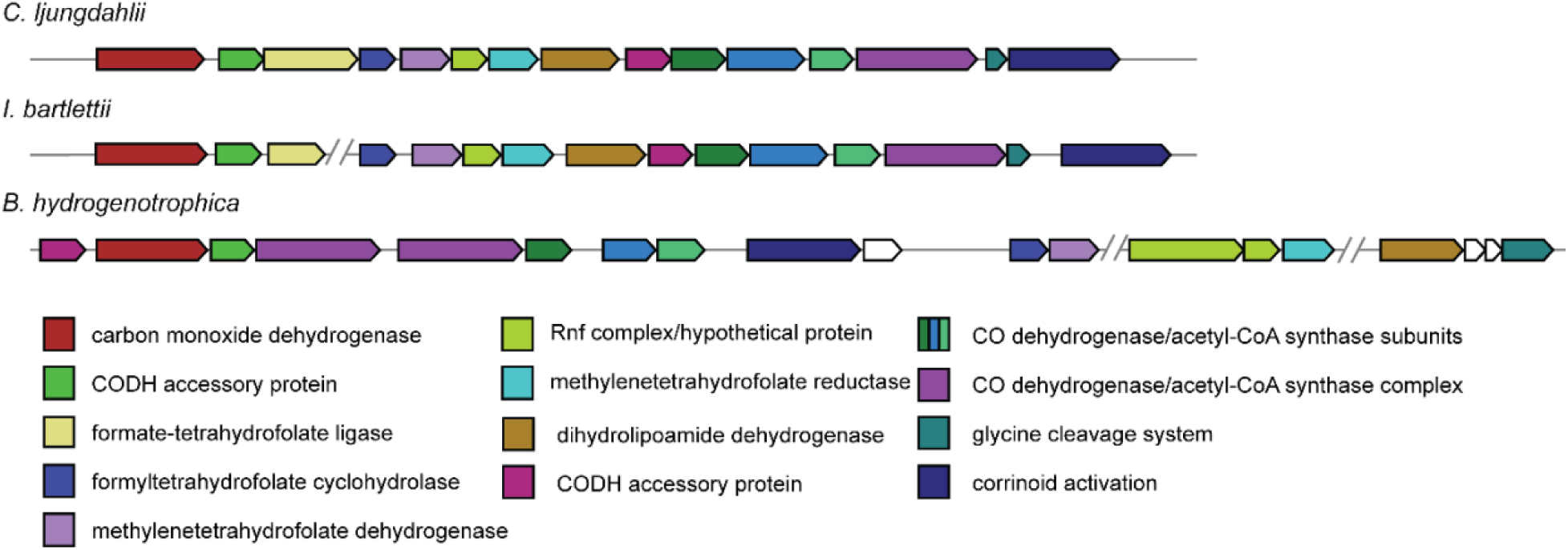
Wood-Ljungdahl Pathway gene clusters in *Iba* and *Bhy*. The *C. ljungdahlii* WLP gene cluster was used as a query to search the genome sequences of *Iba* and *Bhy* using MultiGeneBlast using default parameters.

## Notes

### Competing Interest Statement

The authors have declared no competing interest.

